# A Comprehensive Bibliometric Investigation on Antimicrobials from Fungal Origins with a Biotechnological Perspective

**DOI:** 10.1101/2024.11.18.624177

**Authors:** Lucas da Silva Lopes, Kamila Alves Silva, Thaís da Silva Correa, Mariana Campos-da-Paz, José Guilherme Prado Martin, Ana Alice Maia Gonçalves, Vinícius Silva Belo, Tiago Silveira Gontijo, Alexsandro Sobreira Galdino

## Abstract

Interest in research related to fungi exploration past few years, given its relevance to bioactive compounds production as these metabolites are a promising source of bacteriostatic or fungistatic compounds. Therefore, their use represents a significant alternative to traditional antibiotics by minimizing the risks related to microbial resistance. In this context, this work aimed to: assess the volume of annual publications on the subject and identify key players, analyze the collaboration network among researchers, and check the patents filed on this topic. For this purpose, the Bibliometrix R-package was used, as well as scientific metadata from the Web of Science and Scopus databases (n=506). In total, 256 sources, authors (n=2,526), keywords (n=1,812), and references (n=19,315) were analyzed from 1989 to 2023. The academic debate on the subject has been promoted by India (29%), United Kingdom (UK) (7%), China (6%) and the United States of America (USA) (6%). The authors identified as the most cited were Liu J (n=142), followed by Jesu Arockiaraj (n=106). There is a knowledge predominance of publications focusing on the life science disciplines. The most prolific institutions were the National Research Center (n=20) and the University of Pittsburgh (n=13). The most cited journals were the World Journal of Microbiology & Biotechnology (n=719) and Applied Microbiology and Biotechnology (n=661). Finally, the United States Patent and Trademark Office represented 85% of the patents filed on the subject (n=28,303). Together, these results can direct researchers and Biotech industries in their search for the most important sources related to antimicrobial biotechnology.

## 1. INTRODUCTION

The use of fungi by humans traces back to prehistoric times. It is noteworthy that fungi, as living microorganisms, are considered the primary sources of secondary metabolites, including phenols, steroids, xanthones, peptides, lipids, glycosides, and isoprenoids (Lopes et al., 2011; Fadiji & Babalola, 2020). Nowadays, these metabolites are recognized as potential alternatives for applications in the food, pharmaceutical, and cosmetic industries. A notable example is the industrial production of penicillin by *Penicillium notatum*, described in 1928 by Alexander Fleming; its discovery resulted in a revolution in the medical and pharmaceutical fields (Fleming, 1929). Since then, the synthesis of antibiotics has experienced a significant growth, driven by the discovery of antifungal activity, chemotaxonomy, and fungi chemodiversity (Hussain et al., 2017), prompting the exploration of naturally produced fungal compounds (Cragg et al., 2012). In this sense, the fungi kingdom represents a group of organisms with high value for biotechnological applications (Paul & Joshi, 2022).

Despite these promising applications, the wholesale synthesis and use of antibiotics in the medical and veterinary fields have generated a significant amount of selective pressure, which has resulted in antimicrobial resistance (AMR) development (Mokrani S, Nabti E 2020). AMR is defined as the ability to resist antimicrobials used in the treatment of bacterial, viral, and fungal infections, thereby increasing the risk of disease progression and increased severity (Li, De Oliveira & Walker, 2022). If not adequately controlled, it is expected to result in a loss of $1 billion by 2050, and losses of $1 billion to $3.4 billion in gross domestic product per year by 2030 (Who, 2023, Word Bank, 2017). Additionally, AMR also poses a serious threat to public health, potentially leading to a resurgence of once-treatable infections, longer hospital stays, and rising mortality rates worldwide (Murray et al., 2022). In that regard, considering AMR and issues associated with the toxicity and undesirable side effects of drugs available in the market, the detection of new compounds for microbial control becomes imperative (Sanchez Armengol, Harmanci & Laffleur, 2021).

In the past few decades, several studies on microorganisms as a source of novel antimicrobial targets have been developed (Stierle & Strobel 1993, Strobel & Daisy 2003, Strobel 2003, El-Elimat 2013) However, systematic measurements of studies focused on antimicrobial fungi are scarce, with just a few bibliometric research efforts on essential oil antimicrobials (Ahmad, 2021) as well as the biodiversity of secondary metabolites from bacteria (Al-Shaibani, 2021).

A systematic bibliometric review addresses this gap by providing updates on recent developments, identifying authors involved in central areas, using network analysis to examine citation patterns, and offering bibliographic links to highlight recent advances in the field. Furthermore, this study makes an important contribution to research on antimicrobials of fungal origin by analyzing not only scientific publications, but also the production of patents on the topic and main areas of knowledge, including emerging areas of study, and offering deeper insights into the practical implications and advances in this field.

## 2. METHODS

### 2.1 Data collection

The metadata for this study were selected through a topic search (title, keywords, and abstract) in major scientific repositories relevant to the field: Web of Science (WoS) and Scopus. To conduct a comprehensive survey of publications, records spanning from 1989 (the date of the first publication on the topic) to 2023 (the most recent year with complete information available) were chosen. The descriptors used in the search were “fungal” AND “antimicrobial” AND “biotechnology”. The publications were extracted on February 20, 2024, with conference reports, review articles, books, book chapters, and procedural documents being excluded. The primary reason for exclusion was the absence of research results, such as studies conducted in laboratory settings, and clinical investigations. Additionally, duplicates were removed using R software (R CORE TEAM, 2024), as different search databases might measure identical articles. After reviewing titles and abstracts, records that did not address the topic were excluded, resulting in the elimination of 516 articles. Ultimately, a total of 509 eligible documents were included in the study. The Preferred Reporting Items for Systematic Review and Meta-Analysis Protocols (PRISMA) flowchart was used to graphically represent the process (Figure 1).

**Figure 1.**
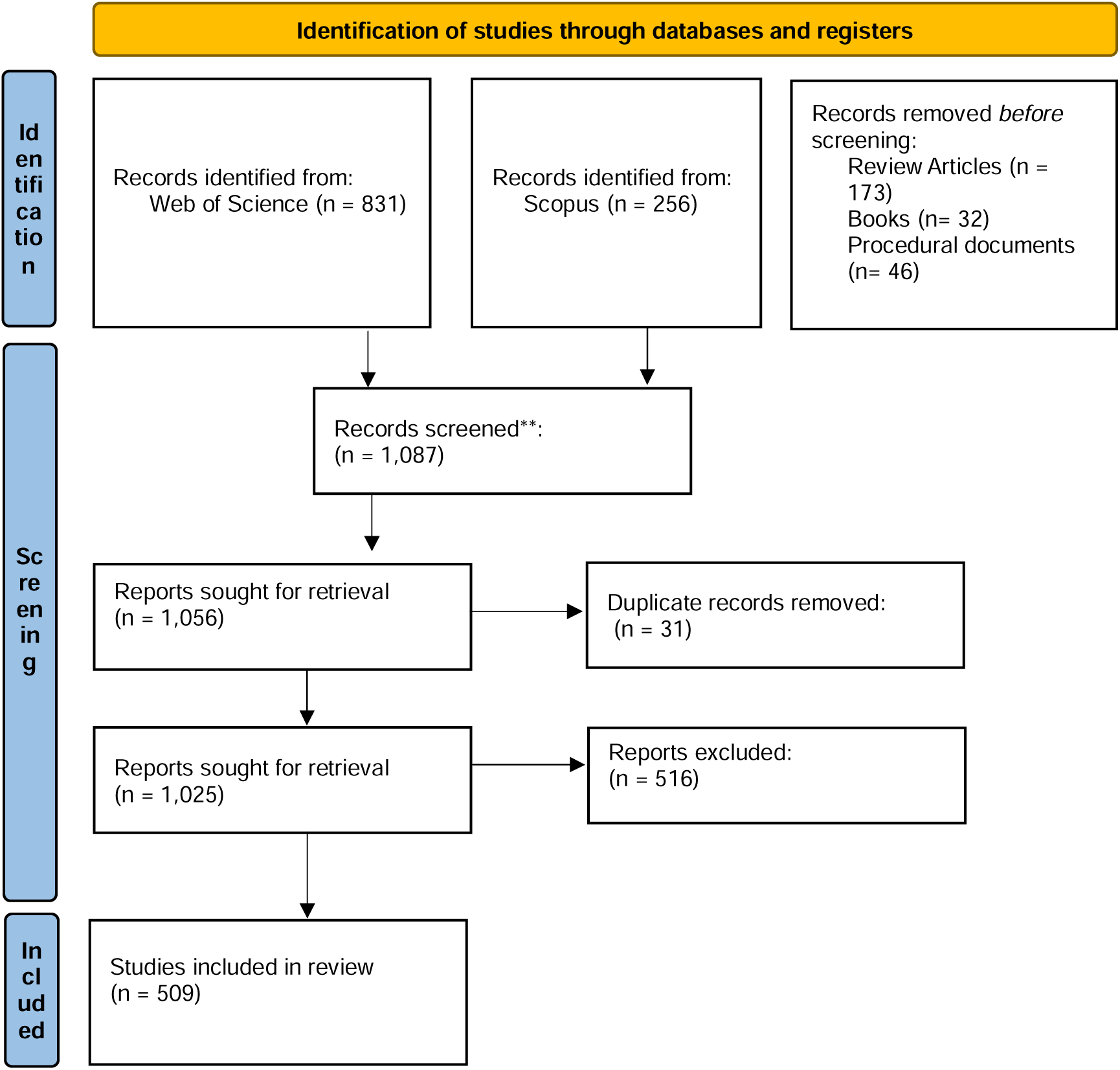
PRISMA Flowchart (http://www.prisma-statement.org/).

Bibliometric analysis, a quantitative method used to capture author networks and keyword occurrences of the research results, was conducted using the R-package Bibliometrix (Aria, M. & Cuccurullo, 2017). By investigating research indicators, it is possible to obtain an overview of the academic characteristics and dynamics, emphasizing strengths and weaknesses that characterize a specific scientific field (Schaer, 2013). This package uses metadata from repository citations to calculate, measure, rank, and identify journal sources. Each article was collected, considering authors, year of publication, title, journal, author affiliations, country, title, and keywords. The data were organized into tables and graphs to facilitate an analysis of the results.

An investigation was also conducted to analyze patents filed on antimicrobials from fungal sources. The Scopus repository was used for this purpose considering the records from 5 different patent offices: World Intellectual Property Organization, European Patent Office, United States Patent and Trademark Office, Japanese Patent Office, and the Intellectual Property Office from the United Kingdom (Kim & Lee, 2015). The patent research was deemed important, as it highlighted potential inventions and encouraged the economic and technological development of new products. The research methods used were selected through a search for topics (title, word, key, and abstract) in the scientific repository Scopus in the patents tool, and records were selected covering the period from 1989 to 2023. The descriptors used in this research were “fungal” AND “antimicrobial” AND “biotechnology”. The publications extracted totaled more than 33,000 patents registered on the platform and to maximize the search, the number of patent writers around the world, patent production, and the relationship between the number of publications over the years and the five most relevant registered patents covering the subject were provided directly by Scopus.

## 3. RESULTS

### 3.1 Annual publications and main indicators

Of the 509 eligible articles, the trends in publications over the years were analyzed. An evolution was observed in the annual growth rate of publications starting with one publication in 1989 and reaching 50 publications in 2023. The Annual Growth Rate of publications for the analyzed period was equal to 12.19%. It reinforced the strength and applicability of antimicrobials from fungal sources over time. The most prolific year in terms of publications was 2020 (n=54) (Figure 2a). Springer (n=29%) was the predominant editorial group, followed by Elsevier (n=17%), and others (n=40%) (Figure 2b). Most of the published articles (n=154) had up to 4 citations, which corroborated the principles of Lotka’s law (Pao, 1985) (Figure 2c). Finally, it appears that most of the published works (n=414) are related to Biotechnology Applied Microbiology (Figure 2d).

**Figure 2:**
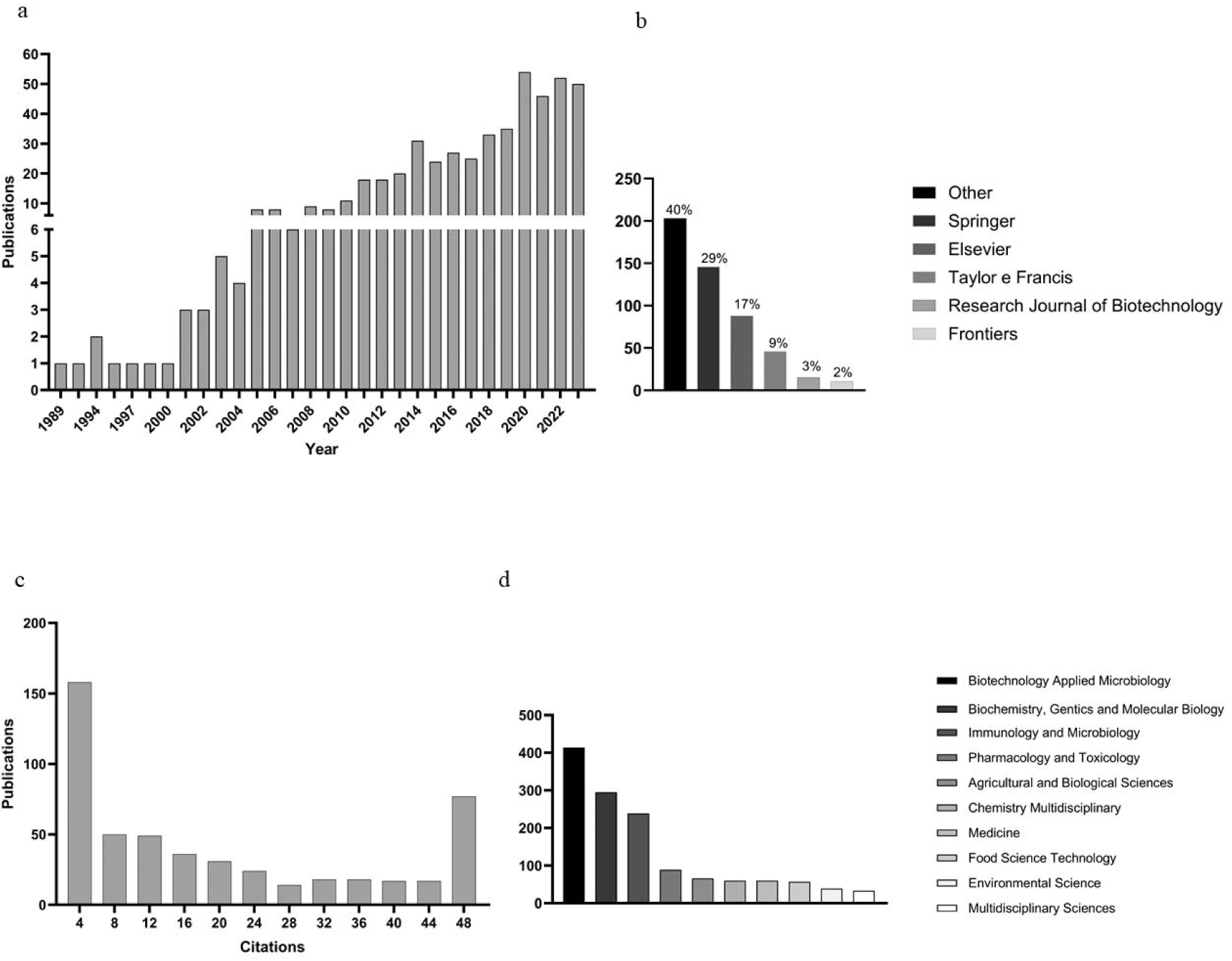
Main information about the publications. Graph (a) shows the number of publications in relation to the year of publication, ranging from 1989 to 2023, as well as the percentage of accumulation over the years (blue line); Graph (b) shows the main publishers in this field, as well as the number of articles published and their degree of percentage; Graph (c) presents the number of articles that have certain citation numbers (ranging from 4 to 4), with the accumulated total highlighted by the orange line; and Graph (d) shows the top 10 areas of knowledge in relation to the number of articles in which they are involved, such as Biotechnology Applied Microbiology, Biochemistry, Genetics and Molecular Biology, Immunology and Microbiology, Pharmacology, Toxicology, and Pharmaceutics.

### 3.2 Geographic distribution of publications

In total, 74 countries have conducted research on fungal-derived antimicrobials over the decades analyzed. The most active contributors came from India, with a frequency of 476 publications, followed by the United States of America (n=162), the United Kingdom (n=162), China (n=142), and Brazil (n = 88) (Figure 3). When analyzing the total number of citations (TC) and the average number of citations per article, India, the UK, and the USA also stood out, with a TC variation ranging from 910 to 3,273, with an average citation between these countries ranging from 22.00 to 27.60 (Table 1).

**Figure 3.**
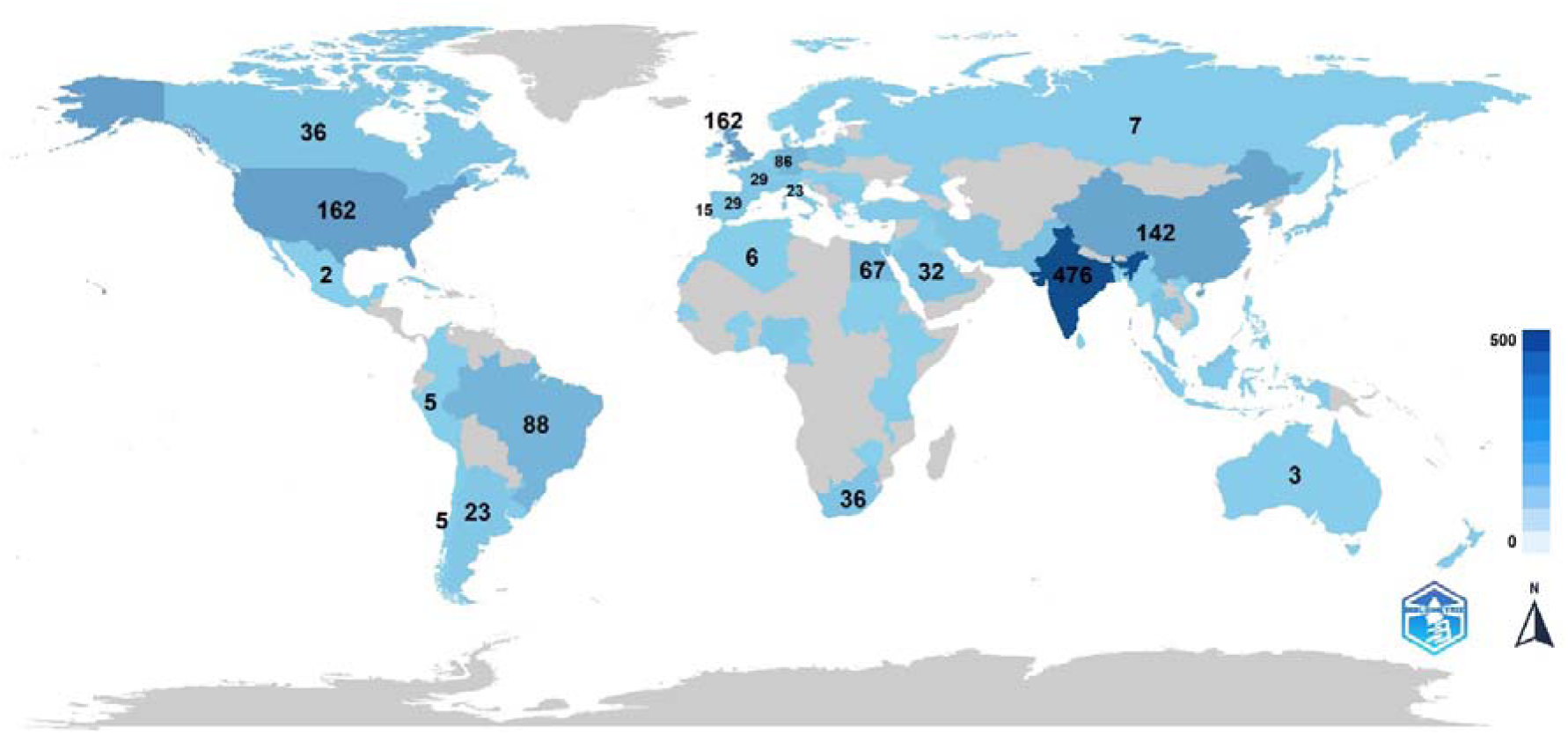
Global Mapping of Publications. Countries with darker shades of blue have higher publication frequencies, while lighter shades denote lower publication rates.

**Table 1.**
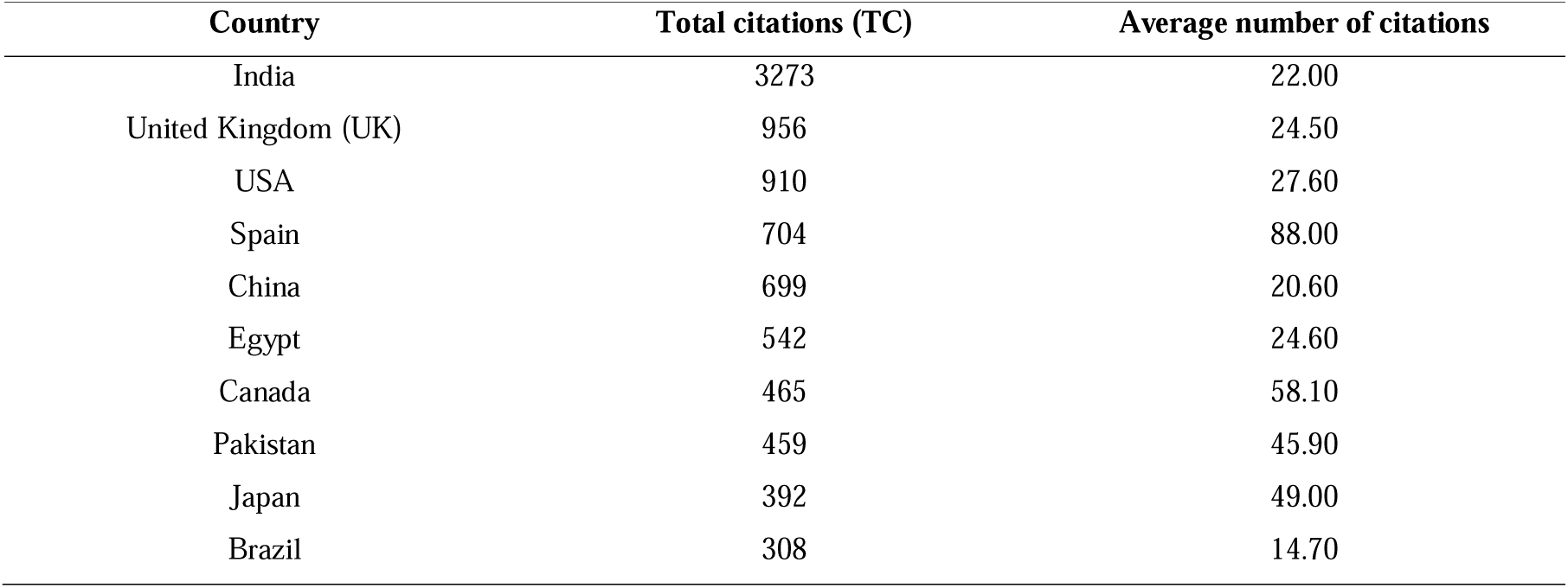
Top Ten Most Cited Countries, 1989–2023.

It should be noted that Spain had the highest average number of citations as compared to any other country, with 88.00, higher than the ranking of the top three countries, demonstrating its worldwide representativeness on the subject (Table 1).

### 3.3 Main research institutions and journals

Table 2 lists the main journals, publisher, impact factor (IF), their h and g indexes, total number of citations (TC), number of publications (NP), and start year. The World Journal of Microbiology & Biotechnology presented the most cited articles, classified by total citations of 719, with an IF of 4.1. Taking these definitions into account, Applied Microbiology and Biotechnology also stood out with indices h and g, respectively, with 13 and 24, followed by Applied Biochemistry and Biotechnology, with a total citation of 661, and an IF of about 5.455.

**Table 2.**
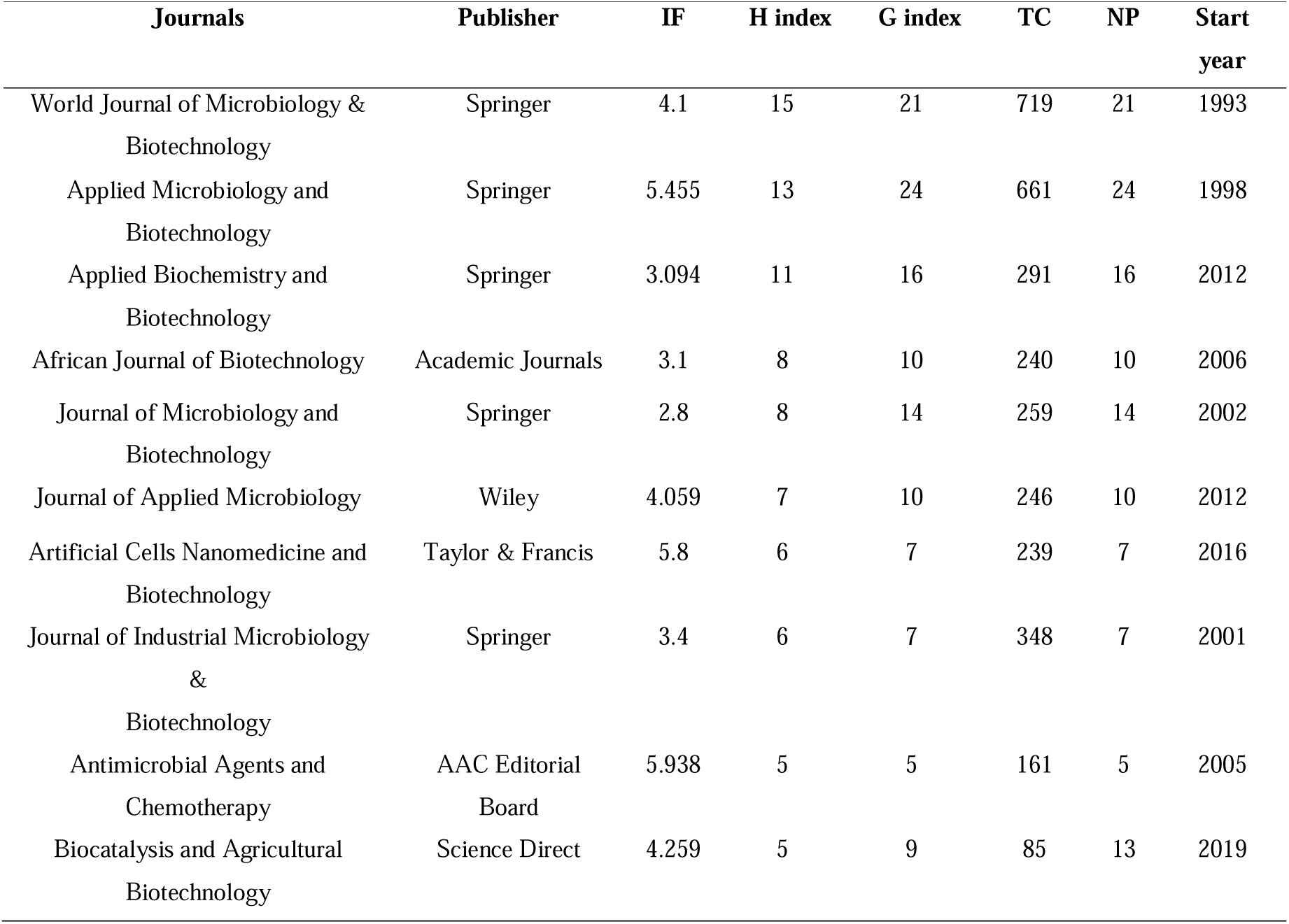
Top 10 Sources and Journals with the Highest Impact.

The key research institutions that made significant contributions to the study are presented in table 3. The National Research Centre, from Egypt, contributed with 20 articles, The University of Pittsburgh, from the United States, with 13 articles, and Universiti Putra Malaysia, from Malaysia, also with 13 articles (Table 3).

**Table 3.**
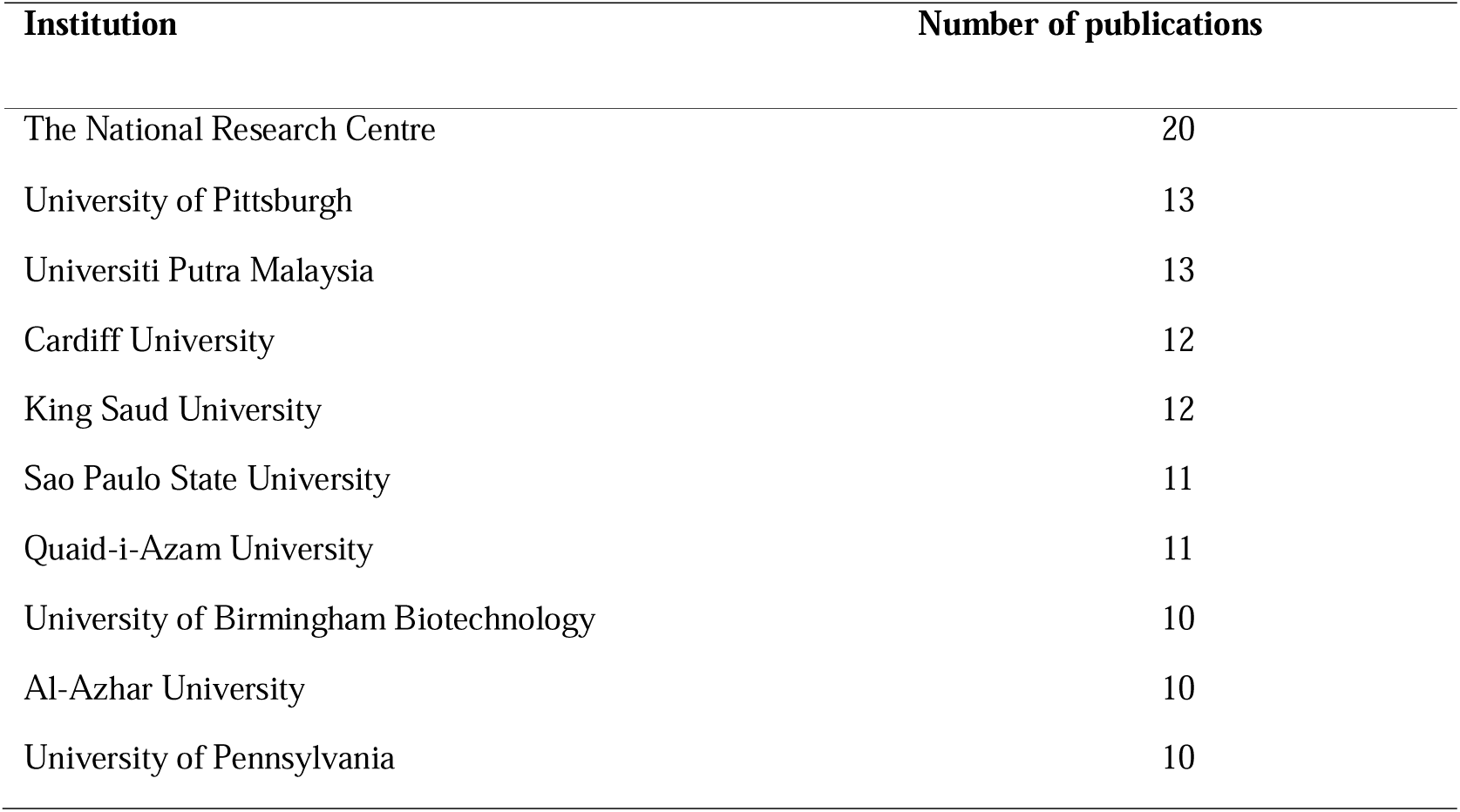
Top 10 Institutions involved in research article production.

### 3.3 Collaboration network and authors

The collaboration network was analyzed through the clusters generated by bibliometrix and their correlation among the authors. An example of this relationship is the greater degree of collaboration and correlation between authors from the same country, such as the author Ya Hao (Yellow Custer) (China), thus strengthening his collaboration networks with authors from the same country. Another example is that of the authors Sira Lee, Juhee Lee, and Seong Ryul Kim (cluster, in orange) (South Korea), whose collaborations are related to lines of research focused on biomolecules and their country of origin, leading to an intimate relationship between the line of research, location, and research institution for the formation of clusters and collaboration networks (Figure 4).

**Figure 4.**
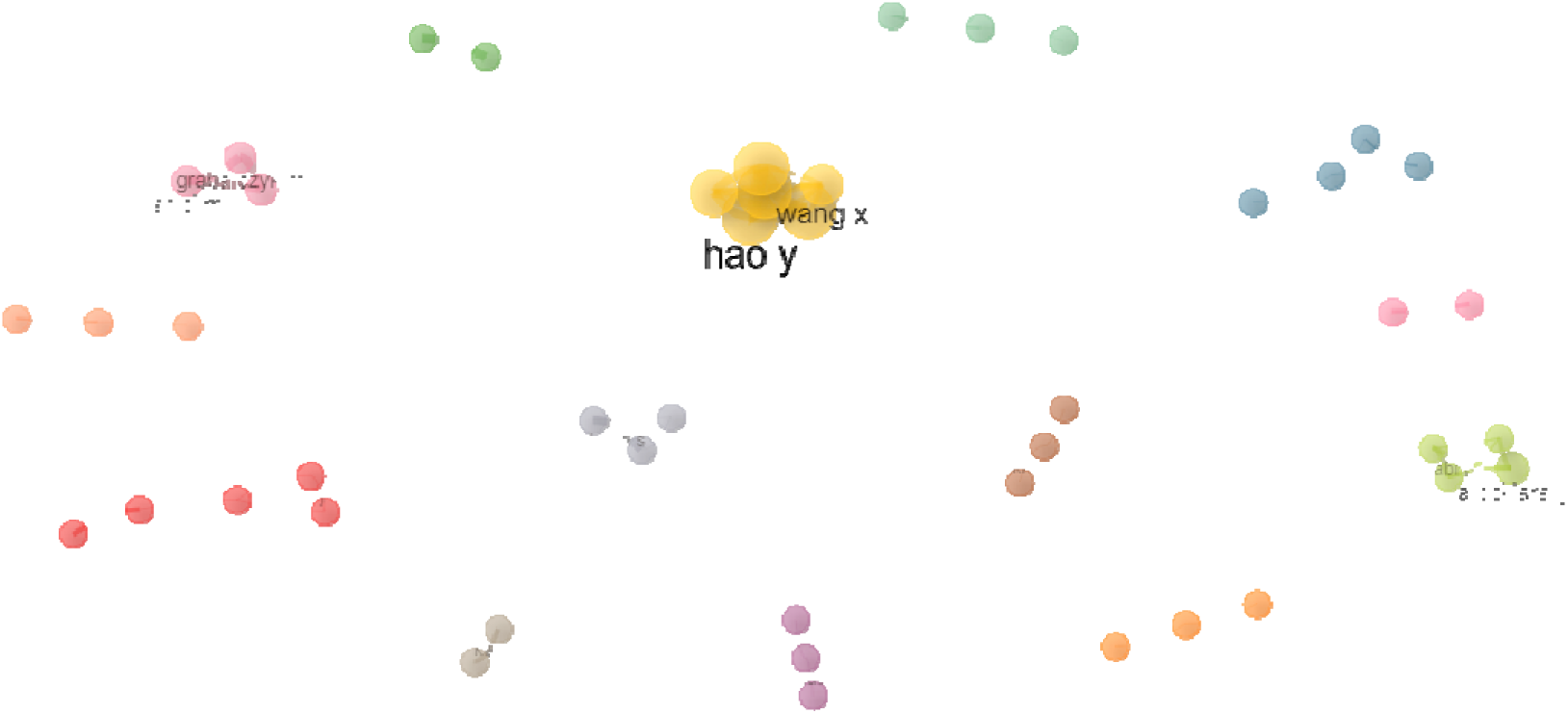
Collaboration Network among Authors. Each cluster, which is defined by a set of terminology used to demonstrate a grouping by similarity that is separated by colors, taking into context groupings of authors by having joint research activity, institution, or country of origin, thus creating a similarity between them.

It is observed that the h indices predominate with values 4, the g indices’ values vary from 4 to 6, the TC values vary from 40 to 143, while the NP values range from 4 to 6. Regarding the authors, their local impact is highlighted by Sira Lee from the Sogang University, South Korea, and Li Y from the Nanjing University,China (Table 4).

**Table 4.**
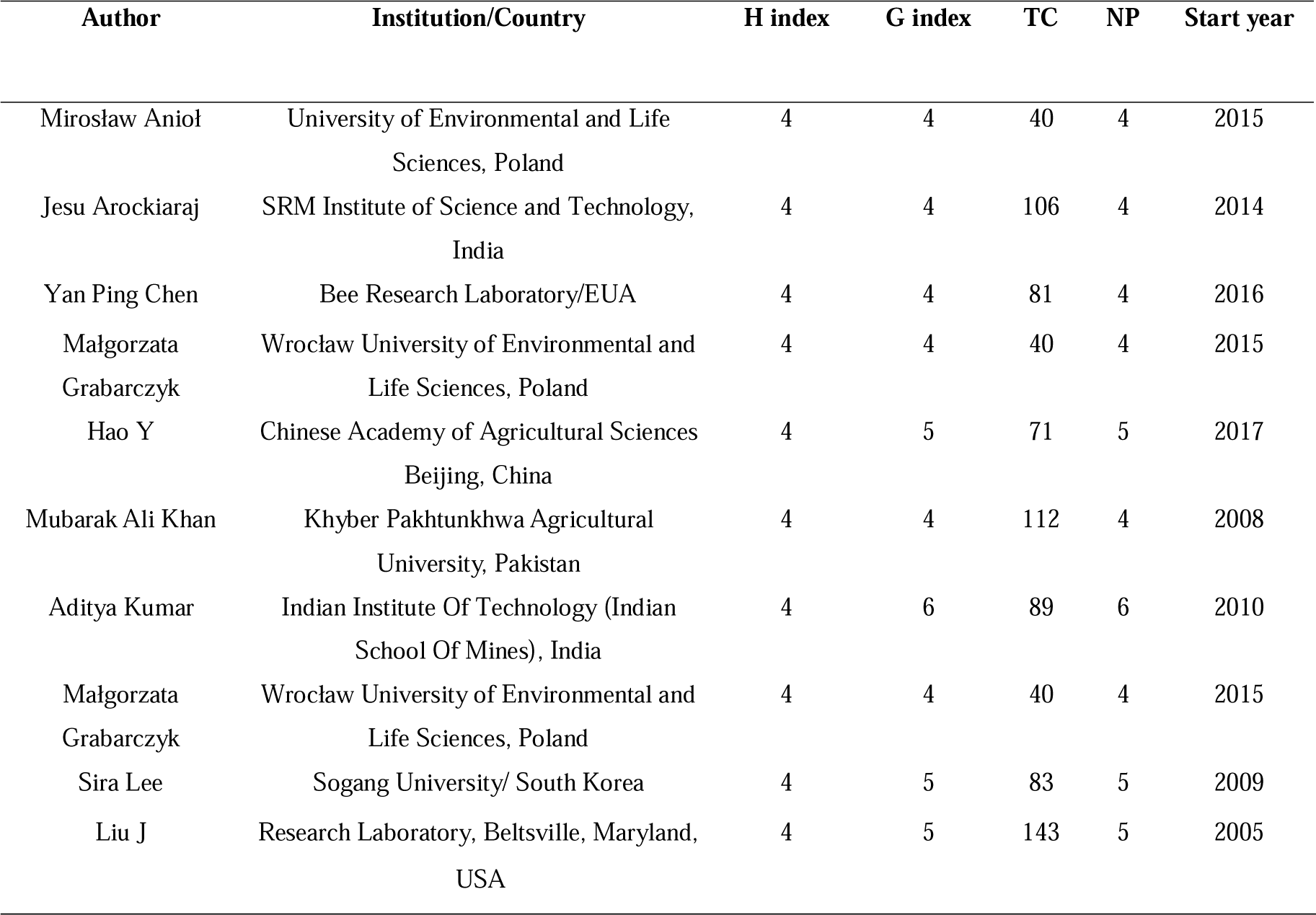
Top 10 Authors and Their Local Impact.

### 3.4 Analysis of co-occurrence of keywords and trending topics

The visualization of Keywords Plus co-occurrence revealed a perspective of reviewed articles to interconnect topics and concepts. Keywords were represented as nodes, and lines functioned as links between the words. The larger the size of a node, the greater the prevalence of a keyword. Additionally, the smaller the distance between the nodes, the more strongly they were related. As expected, “biotechnology” (n=152) “antimicrobial activity” (n=152), and “fungi” (n=139) were the most representative keywords. Furthermore, other keywords such as “nonhuman” (n=136), “fungi“” (n=118), and “article” (n=98) were observed, revealing possible research trends (Figure 5).

**Figure 5.**
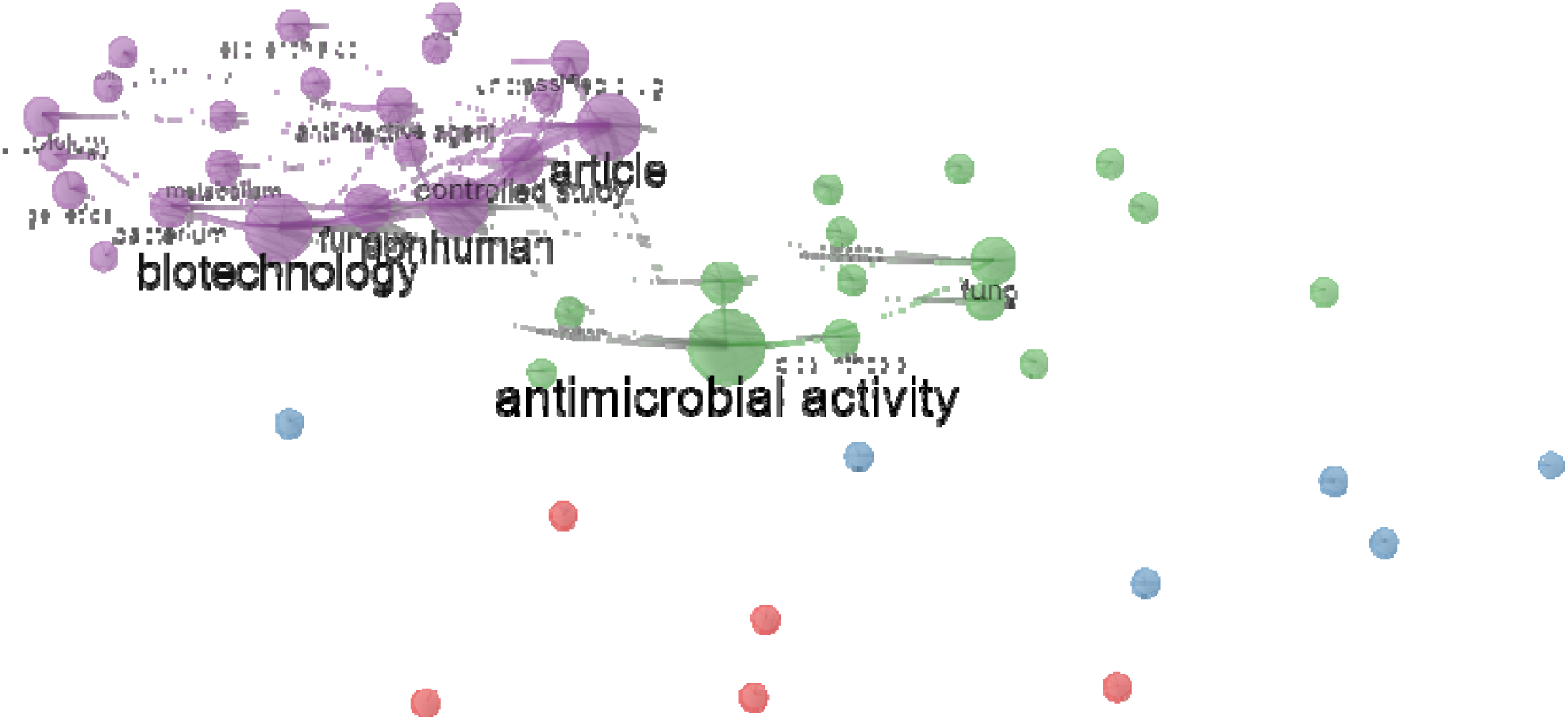
Keyword Co-occurrence Network. Each color represents a subdivision of the broad area of antimicrobials of fungal origin. In green, antimicrobial activities and their branches; in purple, their biotechnological applications; blue represents synthesis and identification methods; and in red, the study of action mechanisms and possible targets of secondary metabolites.

Regarding the most recurring keywords over time, the “trend topics” highlighted the prominent use of “mechanisms” and “pigment” between 2021-2024. Therefore, Figure 6 presents the landscape of the potential emerging areas in antimicrobial application or production.

**Figure 6.**
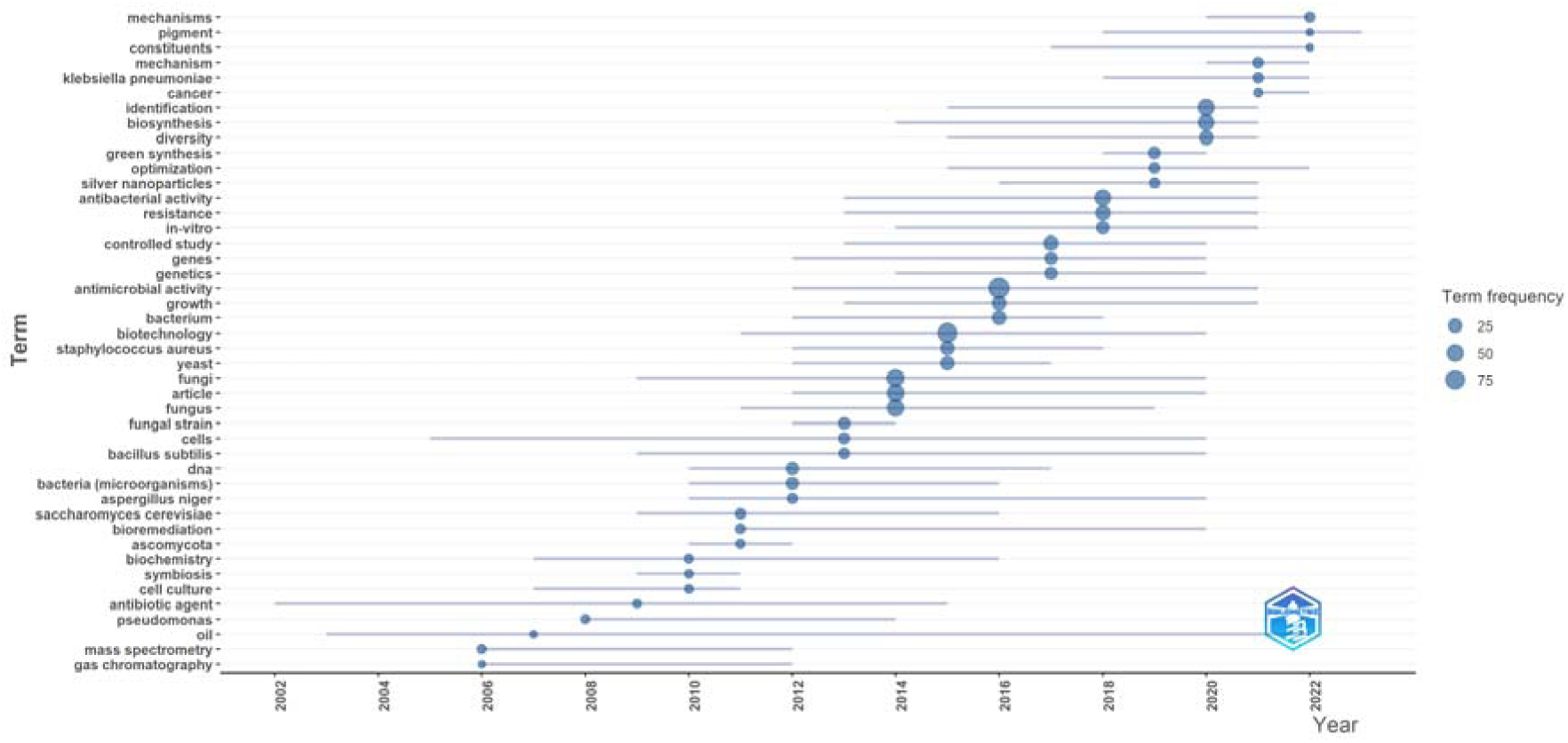
The main highlighted topics on antimicrobial fungi during the period from 2002 to 2024The keyword occurrence ranges from 25 to 75 in which the blue dots vary according to their intensity in relation to the years. The objective of this analysis was not to see the words that appeared most frequently in different studies, but to see the temporal evolution in the use of keywords. The figure in the lower right corner of the graph is the logo for the bibliometrix package.

### 3.5 Analysis of the patent landscape in antimicrobial biotechnology

An analysis of the results obtained in relation to the patent offices found in this study is shown in Figure 8a. The main intellectual property agencies that deal with this subject are: (i) the United States Patent and Trademark Office (www.uspto.gov/) (n = 28,303); (ii) the Japan Patent Office (www.jpo.go.jp/e/) (n=2,599); and (iii) the European Patent Office (www.epo.org/en) (n=2,144). As observed, there has been a substantial growth in the number of patents filed relating to antimicrobial biotechnology, indicating the essential nature of knowledge in this area for various industrial and strategic activities. From 1985 (n=3) to 2023 (n=1,760), 33,397 patents were registered in the Espacenet repository. It is also noteworthy that the patent record occurred in 2021 (n=2,263) (Figure 7b).

**Figure 7:**
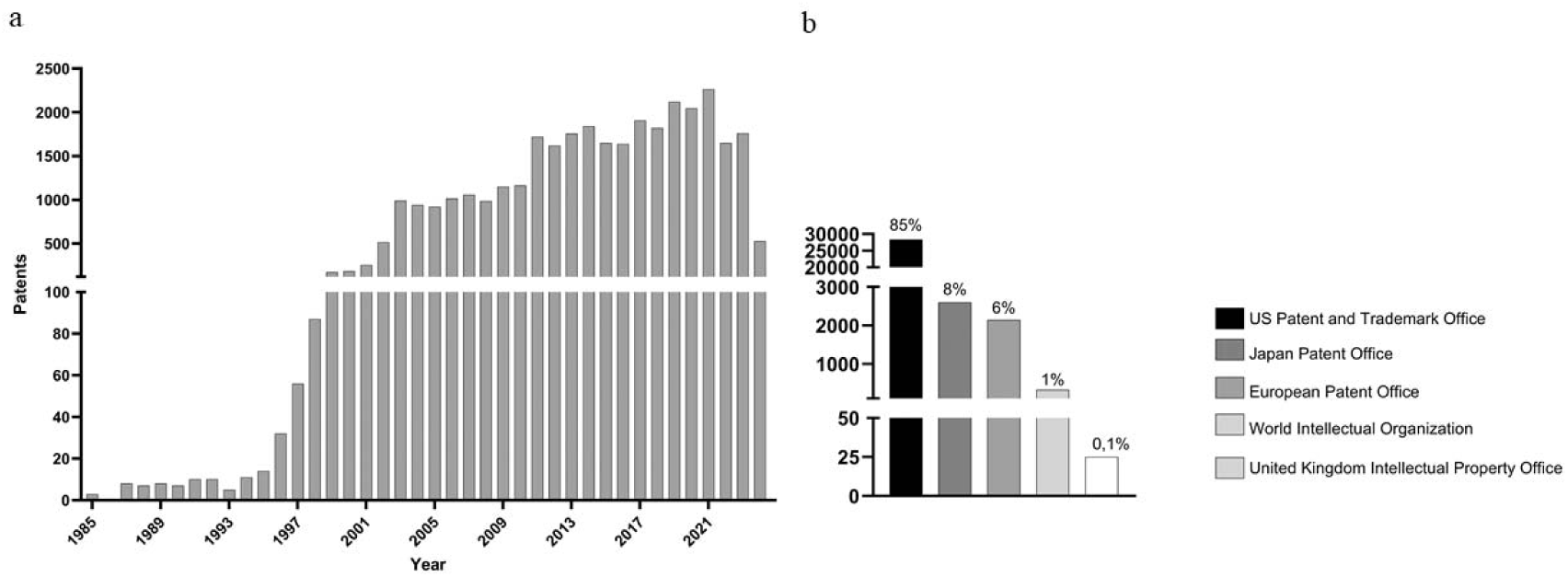
Main information about the patents, 1985 – 2023. (a) number of deposited patents around the world, as well as their relative percentage, and (b) the annual number of patents and a cumulative curve (in blue), showing an exponential trend in patent repositories.

An analysis of the descriptive statistics of the global production of fungal antimicrobial biotechnology patents of the five most relevant patents in the area was also made. The relevance factor was established based on the ranking provided by Espacenet (Table 5). Among the four most relevant patents in the area, there is a hegemony of the United States Patent and Trademark Office, which is evident in its greater weight in patent deposits with 85%, compared to the second-placed European Patent and Trademark Office with only 6%, thus evidencing a very large discrepancy.

**Table 5:**
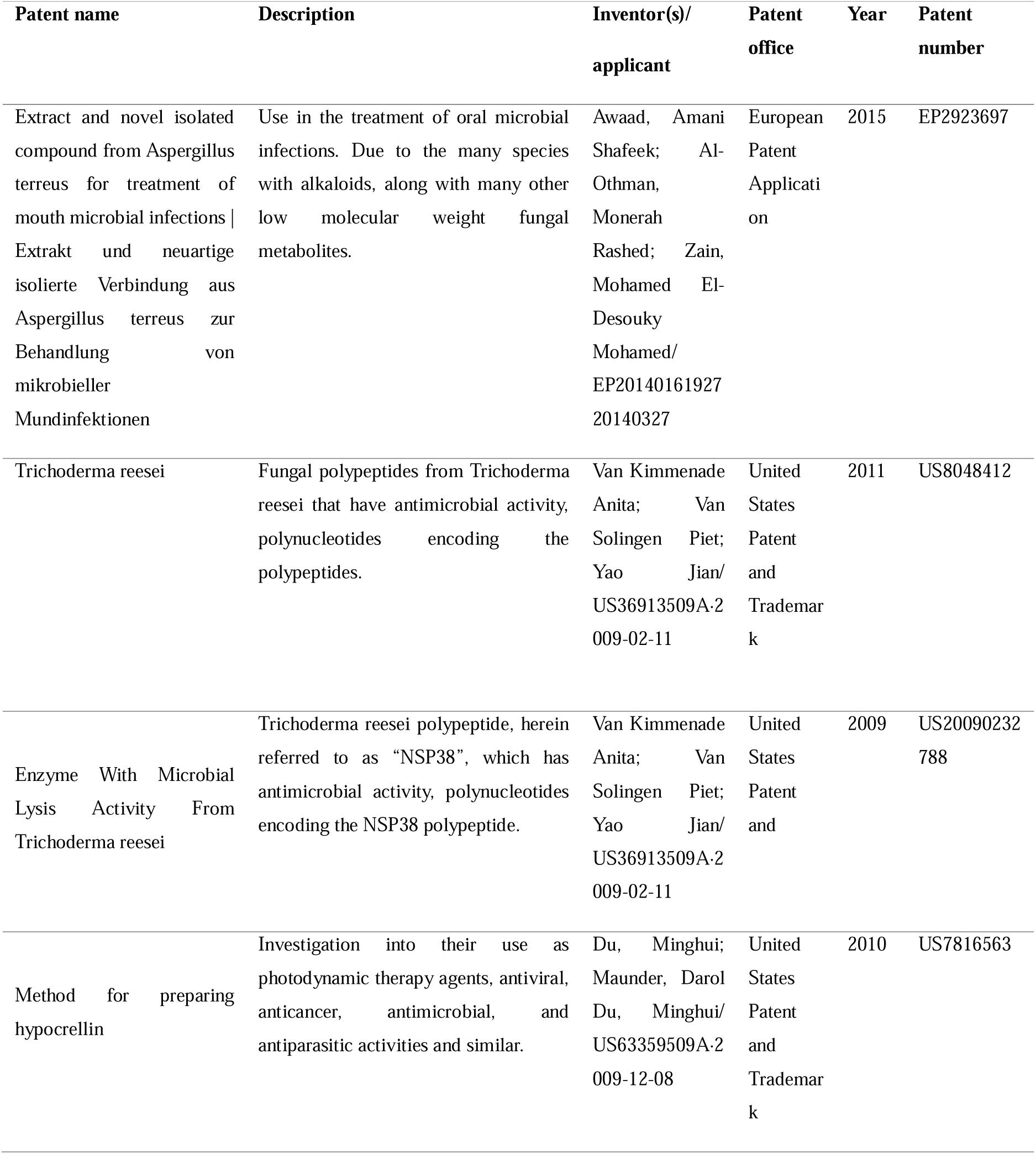
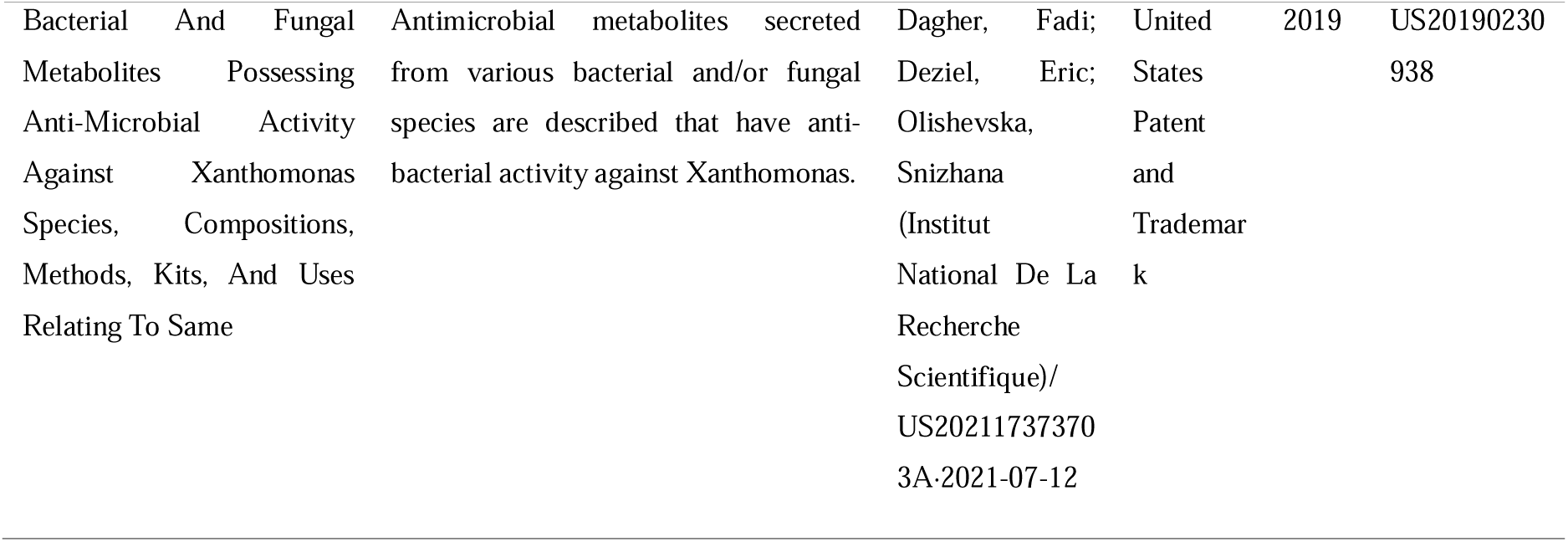
Key patents in terms of relevance and deposit date.

## 4. DISCUSSION

Fungi exhibit a remarkable capacity for biotechnological exploration due to the synthesis of compounds with complex chemical structures, closely related to the biochemical processes of their released substrates (Deacon, 2005). Consequently, fungi are a promising source of bacteriostatic or fungistatic activity against bacterial resistance phenomena, as they produce a broad range of secondary metabolites increasingly sought after to combat antimicrobial resistance (Suay et al., 2000). Therefore, the present investigation collected information through a bibliometric analysis of publications from the WoS and Scopus research repositories. WoS is currently one of the largest databases emphasizing the citation network for prominent articles, a more efficient workflow, and a higher author profile (Pranckutė, 2021). Scopus is a multidisciplinary database with high academic coverage, boasting over 91 million records (Perez-Gilbe, 2023).

The use of the bibliomentrix and biblioshiny interface generates interactive figures that provide information on the disciplinary structure. In this context, the author co-citation networks allow viewing the citations between them individually, as well as the similarity of authors in terms of the frequency with which they are cited by other authors. The co-occurrence of keywords helped the investigation on the similarity of articles that used the same keywords, for instance (Aria & Cuccurullo, 2017; Humboldt-Dachroeden, Rubin & Frid-Nielsen, 2020).

This work revealed that the annual distribution of publications is increasing significantly, due to economic and health problems associated with the increase in infectious and food-borne diseases caused by pathogenic microorganisms. In 2019, 1.27 million deaths worldwide were directly attributed to drug-resistant infections. Around 10 million deaths could occur annually by 2050 (UNEP, 2022). The development of new methodological approaches and the elucidation of the mechanisms of action of secondary metabolites were also revealed, as exemplified in the study conducted by Zanutto et al. (2019).

New methodological approaches to bioprospecting and nanoparticle techniques, as evidenced in the studies by Thokala et al. (2018), are of fundamental importance for strengthening the field of antimicrobials of fungal origin. The main publishers, citations over the years, and areas of knowledge involved in the study were then analyzed. Springer stands out among the publishers, with 146 articles published; in terms of citations, the article with the most publications (n=158) had only 4 citations, although the article with the most publications (n=77) was the most cited. A probable explanation could be the relevance of the subject, different areas, and year of publication. Finally, the analysis of areas of knowledge revealed a predominance of disciplines in life science and health, such Microbial Biotechnology of Microorganisms Biotechnology, Biochemistry, Genetics and Molecular Biology.

Regarding the global mapping of publications, a hegemony was observed between India, the UK, and the USA, being present in almost all the fields that were analyzed in this study. Journal publications and their respective TC were closely linked to their IF. Journals with a higher IF had a lower number of publications (ranging from 5 to 7), such as Antimicrobial Agents and Chemotherapy with 5 publications. Journals with IF ranging from 3.1 to 5.4 had more publications, ranging from 10 and 24, such as Applied Microbiology and Biotechnology and Journal of Applied, respectively. One hypothesis for this result would be the difficulty and costs of publishing in high IF journals, making it inaccessible for authors who end up opting for journals that have low publication costs and consequently low IF. Therefore, funding is of utmost importance so that high-level research can be properly published in high IF journals. In addition, the most prolific institutions on the subject were The National Research/USA Center with 20 publications, followed by the University of Pittsburgh/USA with 13 publications.

The bibliometric analysis also revealed certain gaps, especially about a global and local network of partnerships between countries and authors, with a local prevalence of n=10%, a global prevalence of n=5.67%, and a prevalence of narrower research sub-areas. Through a social network analysis, the author collaboration network was seen to be made up of small author collaboration groups and the authors had affiliated networks in terms of line of research, location, and institution. Such networks revealed many interesting features of academic communities, such as establishing communication networks, sharing ideas, resources, and information, and increasing research productivity (Fonseca, 2016). The importance of strengthening and creating collaborative networks with the aim of disseminating research, exchanging knowledge, and the possibility of researchers being present in different areas of research conducted through these collaborations is evident.

A possible explanation for these recurring keywords could be deeper studies on the molecular identification of biological compounds synthesized by fungi and their antimicrobial action mechanisms driven by the development of techniques for genomic DNA isolation and phylogenetic analysis (Ghaffari F;Ebadi M;Mollaei S. 2023). In one study, Ghaffari et al. (2023) investigated the molecular identification, analysis of fatty acid profiles, and antibacterial properties of endophytic fungi.

Concerning “pigment,” there is currently a high demand for less harmful dyes with biotechnological potential, which is why fungi offer a cheaper alternative with fewer deleterious effects as compared with synthetics (Sousa, 2023). Additionally, many fungi exhibit important antibacterial, antioxidant, and antitumor biological activities and offer a wide range of colors (Kumar et al., 2017, Rao et al., 2017). Sousa et al. (2023) isolated and identified endophytic filamentous fungus, as well as the chemical characterization and evaluation of fluorescence, toxicity, stability, and potential application of its synthesized red dye.

In terms of patent registrations, the United States Patent & Trademark holds 28,303 of the 33,397 patents registered, demonstrating its strength on the world stage. As shown above, there has been an exponential growth in scientific publications in the antimicrobial field, which can be directly related to a massive increase in patent registrations. It should be noted that, due to data availability, these analyses were based on the Espacenet repository. While scientific literature provides theoretical foundations and insights, patents cover applied research into possible innovations. The analysis and discovery of exorbitant patent filings underscored the importance of patenting a product or discovery as it brings transparency and security to researchers and consumers, in addition to guarantees of exclusivity and investment in technology innovation. Furthermore, having the largest number of patents filed does not necessarily mean having the largest number of commercial products. This is because not every patent becomes a product, thereby highlighting the importance of filing patents. It can transfer technology to the productive sector because, although patents do not become products, they burden universities and research centers with annual maintainence fees until they are granted.

The study’s limitations relate to the discrepancy between the results obtained in different search databases and the distinction in search topics between both. The initial screening revealed a total search on the WoS of (n=819) and on Scopus of (n=252), with a difference interval of about 567 articles Therefore, changes were made to both the keywords and topics to be searched and the Boolean operators. However, this resulted in either a very specific number with few articles or a very comprehensive one. Therefore, to reach an acceptable number that covered a plausible number of articles, such changes were made. Research on fungal antimicrobial biotechnology holds remarkable theoretical significance. However, there is a gap to be filled for future studies to not only examine intellectual output, but also scrutinize technological products and patents generated in this domain. Such an approach will enable a more comprehensive overview, offering deeper insights into the practical implications and advances within this field.

## 5. CONCLUSIONS

This study presented a more comprehensive approach to aspects of annual publications, geographic distributions, the most published journals, the most prolific authors, emerging areas of study, main areas of knowledge, perspectives on patents, and offers deeper insights into practical implications and advances in this field. Regarding future perspectives, bioinformatics and synthetic biology will allow significant advances with the use of heterologous gene expression, sequences, and mechanisms of action of proteins and in silico antimicrobial peptides that will make it possible to synthesize metabolites on a large scale and apply them.

## Supporting information

Tables of Manuscript

## LIST OF ABBREVIATIONS

(AMR): Antimicrobial resistance
(IF): Impact factor
(TC): Total number of citations
(PRISMA): Systematic Review and Meta-Analysis Protocols
(NP): Number of publications
(WoS): Web of Science

## ACKNOWLEDGEMENTS

The authors would like to thank CAPES (Finance Code 001), CNPq, and FAPEMIG for the scholarships. RCG, EAFC and ASG would like to thank CNPq for their research fellowship. The authors are grateful to UFSJ, UFV, CAPES, CNPq and FAPEMIG.

