## Supplementary material for "A Comprehensive Bibliometric Investigation on Antimicrobials from Fungal Origins with a Biotechnological Perspective": Tables of Manuscript

Table 1. Top Ten Most Cited Countries, 1989–2023.

| **Country** | **Total citations (TC)** | **Average number of citations** |
| --- | --- | --- |
| India | 3273 | 22.00 |
| United Kingdom (UK) | 956 | 24.50 |
| USA | 910 | 27.60 |
| Spain | 704 | 88.00 |
| China | 699 | 20.60 |
| Egypt | 542 | 24.60 |
| Canada | 465 | 58.10 |
| Pakistan | 459 | 45.90 |
| Japan | 392 | 49.00 |
| Brazil | 308 | 14.70 |

Table 2. Top 10 Sources and Journals with the Highest Impact.

| **Journals** | **Publisher** | **IF** | **H index** | | **G index** | **TC** | **NP** | **Start year** |
| --- | --- | --- | --- | --- | --- | --- | --- | --- |
| World Journal of Microbiology & Biotechnology | Springer | 4.1 | 15 | 21 | | 719 | 21 | 1993 |
| Applied Microbiology and Biotechnology | Springer | 5.455 | 13 | 24 | | 661 | 24 | 1998 |
| Applied Biochemistry and Biotechnology | Springer | 3.094 | 11 | 16 | | 291 | 16 | 2012 |
| African Journal of Biotechnology | Academic Journals | 3.1 | 8 | 10 | | 240 | 10 | 2006 |
| Journal of Microbiology and Biotechnology | Springer | 2.8 | 8 | 14 | | 259 | 14 | 2002 |
| Journal of Applied Microbiology | Wiley | 4.059 | 7 | 10 | | 246 | 10 | 2012 |
| Artificial Cells Nanomedicine and Biotechnology | Taylor & Francis | 5.8 | 6 | 7 | | 239 | 7 | 2016 |
| Journal of Industrial Microbiology &  Biotechnology | Springer | 3.4 | 6 | 7 | | 348 | 7 | 2001 |
| Antimicrobial Agents and Chemotherapy | AAC Editorial Board | 5.938 | 5 | 5 | | 161 | 5 | 2005 |
| Biocatalysis and Agricultural Biotechnology | Science Direct | 4.259 | 5 | 9 | | 85 | 13 | 2019 |

Table 3. Top 10 Institutions involved in research article production

| **Institution** | **Number of publications** |
| --- | --- |
| The National Research Centre | 20 |
| University of Pittsburgh | 13 |
| Universiti Putra Malaysia | 13 |
| Cardiff University | 12 |
| King Saud University | 12 |
| Sao Paulo State University | 11 |
| Quaid-i-Azam University | 11 |
| University of Birmingham Biotechnology | 10 |
| Al-Azhar University | 10 |
| University of Pennsylvania | 10 |

Table 4. Top 10 Authors and Their Local Impact

| **Author** | **Institution/Country** | **H index** | **G index** | **TC** | **NP** | **Start year** |
| --- | --- | --- | --- | --- | --- | --- |
| Mirosław Anioł | University of Environmental and Life Sciences, Poland | 4 | 4 | 40 | 4 | 2015 |
| Jesu Arockiaraj | SRM Institute of Science and Technology, India | 4 | 4 | 106 | 4 | 2014 |
| Yan Ping Chen | Bee Research Laboratory/EUA | 4 | 4 | 81 | 4 | 2016 |
| Małgorzata Grabarczyk | Wrocław University of Environmental and Life Sciences, Poland | 4 | 4 | 40 | 4 | 2015 |
| Hao Y | Chinese Academy of Agricultural Sciences Beijing, China | 4 | 5 | 71 | 5 | 2017 |
| Mubarak Ali Khan | Khyber Pakhtunkhwa Agricultural University, Pakistan | 4 | 4 | 112 | 4 | 2008 |
| Aditya Kumar | Indian Institute Of Technology (Indian School Of Mines), India | 4 | 6 | 89 | 6 | 2010 |
| Małgorzata Grabarczyk | Wrocław University of Environmental and Life Sciences, Poland | 4 | 4 | 40 | 4 | 2015 |
| Sira Lee | Sogang University/ South Korea | 4 | 5 | 83 | 5 | 2009 |
| Liu J | Research Laboratory, Beltsville, Maryland, USA | 4 | 5 | 143 | 5 | 2005 |

Table 5: Key patents in terms of relevance and deposit date

| Patent name | Description | Inventor(s)/  applicant | Patent office | Year | Patent number |
| --- | --- | --- | --- | --- | --- |
| Extract and novel isolated compound from Aspergillus terreus for treatment of mouth microbial infections \| Extrakt und neuartige isolierte Verbindung aus Aspergillus terreus zur Behandlung von mikrobieller Mundinfektionen | Use in the treatment of oral microbial infections. Due to the many species with alkaloids, along with many other low molecular weight fungal metabolites. | Awaad, Amani Shafeek; Al-Othman, Monerah Rashed; Zain, Mohamed El-Desouky Mohamed/ EP20140161927 20140327 | European Patent Application | 2015 | EP2923697 |
| Trichoderma reesei | Fungal polypeptides from Trichoderma reesei that have antimicrobial activity, polynucleotides encoding the polypeptides. | Van Kimmenade Anita; Van Solingen Piet; Yao Jian/ US36913509A·2009-02-11 | United States Patent and Trademark | 2011 | US8048412 |
| Enzyme With Microbial Lysis Activity From Trichoderma reesei | Trichoderma reesei polypeptide, herein referred to as “NSP38”, which has antimicrobial activity, polynucleotides encoding the NSP38 polypeptide. | Van Kimmenade Anita; Van Solingen Piet; Yao Jian/ US36913509A·2009-02-11 | United States Patent and | 2009 | US20090232788 |
| Method for preparing hypocrellin | Investigation into their use as photodynamic therapy agents, antiviral, anticancer, antimicrobial, and antiparasitic activities and similar. | Du, Minghui; Maunder, Darol Du, Minghui/ US63359509A·2009-12-08 | United States Patent and Trademark | 2010 | US7816563 |
| Bacterial And Fungal Metabolites Possessing Anti-Microbial Activity Against Xanthomonas Species, Compositions, Methods, Kits, And Uses Relating To Same | Antimicrobial metabolites secreted from various bacterial and/or fungal species are described that have anti-bacterial activity against Xanthomonas. | Dagher, Fadi; Deziel, Eric; Olishevska, Snizhana (Institut National De La Recherche Scientifique)/ US202117373703A·2021-07-12 | United States Patent and Trademark | 2019 | US20190230938 |
